# Temporal codes of visual working memory in the human cerebral cortex

**DOI:** 10.1101/2020.04.26.062752

**Authors:** Yasuki Noguchi, Ryusuke Kakigi

**Affiliations:** Department of Psychology, Graduate School of Humanities, Kobe University, 1-1 Rokkodai-cho, Nada, Kobe, 657-8501, Japan; Department of Integrative Physiology, National Institute for Physiological Sciences, 38 Nishigonaka, Myodaiji, Okazaki, 444-8585, Japan

**Keywords:** magnetoencephalography, change detection, alpha frequency, beta frequency, front-parietal cortex

## Abstract

Visual working memory (vWM) is an important ability required for various cognitive tasks although its neural underpinnings remain controversial. While many studies have focused on theta (4-7 Hz) and gamma (> 30 Hz) rhythms as a substrate of vWM, here we show that temporal signals embedded in alpha (8-12 Hz) and beta (13-30 Hz) bands can be a good predictor of vWM capacity. Neural activity of healthy human participants was recorded with magnetoencephalography when they performed a classical vWM task (change detection). We analyzed changes in inter-peak intervals (IPIs) of oscillatory signals along with an increase in WM load (a number of to-be-memorized items, 1-6). Results showed a load-dependent reduction of IPIs in the parietal and frontal regions, indicating that alpha/beta rhythms became faster when multiple items were stored in vWM. Furthermore, this reduction in IPIs was positively correlated with individual vWM capacity, especially in the frontal cortex. Those results indicate that vWM is represented as a change in oscillation frequency in the human cerebral cortex.

## Introduction

Working memory (WM) is a fundamental ability in various cognitive skills, such as goal-directed behaviors, inference, calculations, and decision making (Baddeley, 2012; Luck and Vogel, 2013; Ma et al., 2014; D’Esposito and Postle, 2015; Eriksson et al., 2015; Kaminski et al., 2017; Leavitt et al., 2017; Jacob et al., 2018). Although its neural mechanisms remain a matter of debate (Stokes, 2015; Constantinidis et al., 2018; Lundqvist et al., 2018), an increasing number of studies have reported a close relationship of WM with oscillatory signals in the brain (Roux and Uhlhaas, 2014; Kornblith et al., 2016; Johnson et al., 2017). Especially, a model based on the cross-frequency coupling between theta (4-7 Hz) and gamma (> 30 Hz) rhythms has gained much attention (Canolty et al., 2006; Sauseng et al., 2009; Axmacher et al., 2010; Lisman and Jensen, 2013; Miller et al., 2018; Reinhart and Nguyen, 2019).

In contrast to the theta and gamma rhythms, a role of alpha (8-12 Hz) and beta (13-30 Hz) rhythms in WM has been controversial, partly because those rhythms are thought to reflect a suppression of neural processing (Jensen and Mazaheri, 2010; van Ede et al., 2011; Shin et al., 2017). Several studies reported an increase in power of alpha and beta bands during a retention period of vWM tasks (Jensen et al., 2002; Palva et al., 2011). According to Bonnefond & Jensen (2012), this increased alpha activity reflected a suppression of neural processing for sensory inputs during the retention period, which protected WM contents from retroactive interference (Bonnefond and Jensen, 2012). Other studies also indicated a functional (facilitatory) role of alpha/beta activity in WM maintenance (Spitzer and Haegens, 2017; Gelastopoulos et al., 2019; Wianda and Ross, 2019). In contrast, there are some studies reporting a negative (inhibitory) aspect of alpha/beta rhythms in vWM (Fukuda et al., 2015; Proskovec et al., 2018; Mapelli and Ozkurt, 2019). For example, Mapelli & Özkurt (2019) showed that alpha-beta activity in the parieto-occipital cortex was larger in incorrectly-than correctly-answered trials of a vWM task. The enhanced alpha/beta powers in a retention period (Jensen et al., 2002; Palva et al., 2011) therefore might reflect the decay or “clearing-out” (not maintenance) of WM contents (Miller et al., 2018; Schmidt et al., 2019).

Most studies listed above have investigated changes in power (amplitude) of alpha/beta rhythm related to vWM. Here we address this issue by measuring a *speed* of those waves, using the inter-peak interval (IPI) analysis of magnetoencephalography (MEG) (Noguchi et al., 2019; Noguchi and Kubo, 2020). In this analysis, raw waveforms of MEG are filtered with a pass-band of interest (e.g. 8-30 Hz for alpha-to-beta band). Each interval between contiguous peaks in the filtered waveforms is defined as IPI (Fig. 1A). Changes in mean IPI lengths indicate changes in oscillation frequency (speed) of alpha/beta waves (faster rhythm produces shorter IPIs, Fig. 1B). This approach focusing on temporal codes of oscillatory signals might reveal a new role of alpha/beta rhythms in vWM.

**Figure 1.**
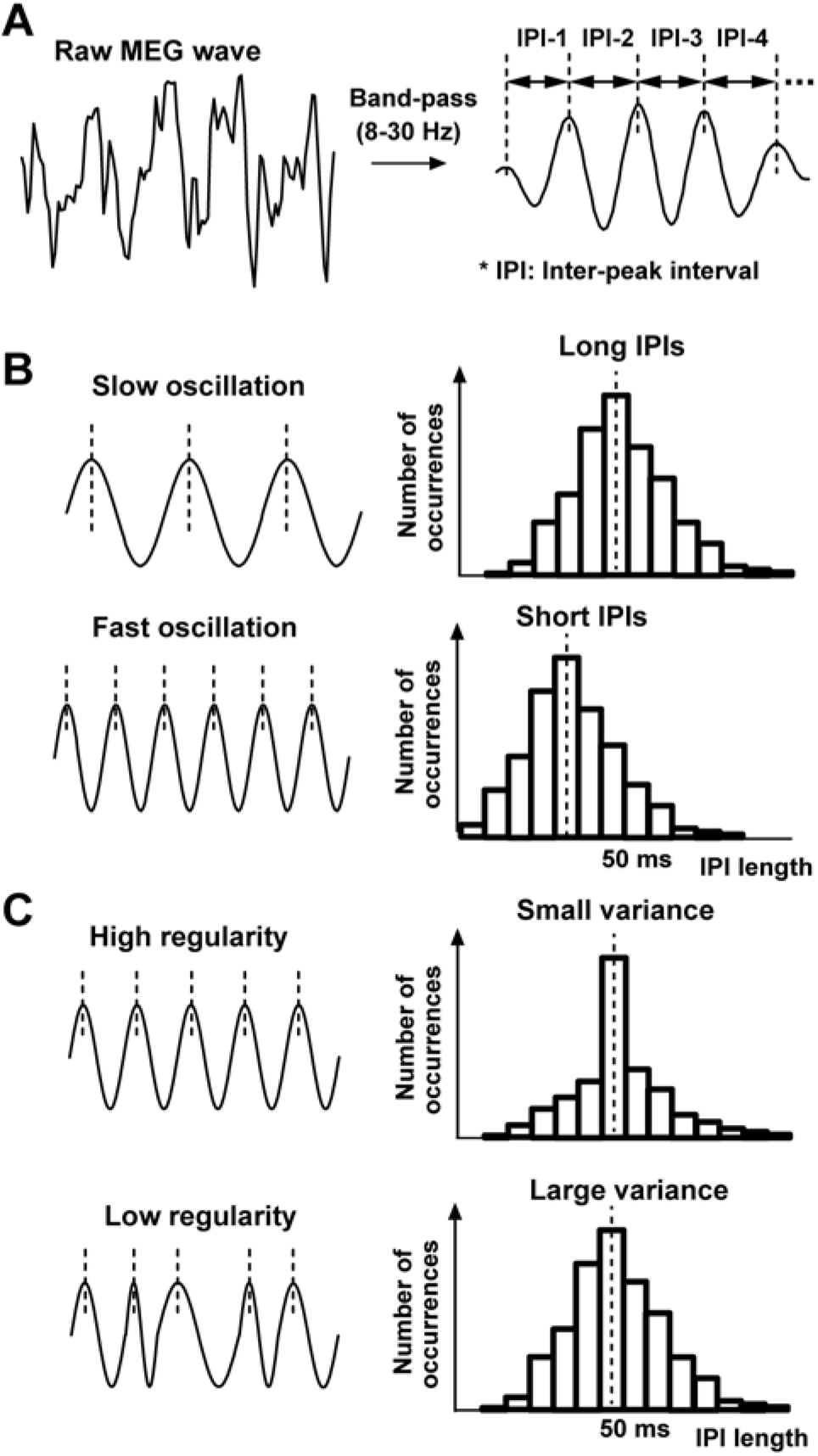
Schematic illustrations of the inter-peak interval (IPI) analysis. We first extracted oscillatory signals at alpha-to-beta band (8–30 Hz) with a band-pass filter (**A**). Each IPI was measured as a time length between two peaks of the filtered waveform. We then pooled all IPIs within a retention period (300 – 900 ms, Fig. 2) across trials, depicting a distribution of their occurrences. Since slow/fast oscillatory signals produce longer/shorter IPIs, changes in oscillation frequency can be measured as changes in mean IPIs (**B**). On the other hand, the regularity of oscillatory signals can be indexed by a variance of the IPI distribution (**C**), because an irregular signal produces many IPIs distant from the mean.

One advantage of our IPI analysis is that it can evaluate not only speed but also the *regularity* of brain rhythms; a smaller variance of an IPI distribution indicates higher regularity of oscillatory signal (Fig. 1C). Here we use this approach to test the “attractor hypothesis” in WM. Although there is intense debate about neural underpinnings of WM, electrophysiological studies on animals have proposed the attractor states (stable fixed points in collective neuronal activity) as a key mechanism for memory retention (Compte et al., 2000; Brody et al., 2003; Chaudhuri and Fiete, 2016; Murray et al., 2017; Inagaki et al., 2019). We expect that those attractor states might be measured as a reduced variance of IPIs, because they emerge from a self-sustained recurrent neural network (Macoveanu et al., 2006; Wimmer et al., 2014; Standage and Pare, 2018) that can generate periodic (regular) signals. Such data would provide further evidence for the attractor hypothesis, elucidating neural substrates of WM in the normal human brain.

## Materials and Methods

### Participants

Twenty-four healthy human subjects (18 females, age: 19-50) with normal or corrected-to-normal vision participated in the present study. This sample size was based on the power analysis using G*Power 3 (Faul et al., 2007). An effect size was estimated from data of our previous MEG study (Noguchi et al., 2019), with the type I error rate (alpha) and statistical power (1 - beta) set at < 0.05 and > 0.80, respectively. Because the data of three participants contained excessive noise in MEG waveforms (due to dental treatments), they were discarded and replaced by thsoe of additional three subjects. Informed consent was received from each participant after the nature of the study had been explained. All experiments were carried out in accordance with regulations and guidelines approved by the ethics committee of Kobe University, Japan.

### Stimuli and task

We measured vWM capacity of the participants using a cued change-detection task (Vogel and Machizawa, 2004). All visual stimuli were generated with the Matlab Psychophysics Toolbox (Brainard, 1997; Pelli, 1997) and projected on a screen at a refresh rate of 60 Hz. Each trial started with a black fixation point (0.18 ×0.18 deg) over a gray background for 800 ms. To direct attention of participants, a cue stimulus (an arrow pointing leftward or rightward, 1.34 deg) was presented over the central field for 33 ms (Fig. 2A), followed by another fixation period of 467 ms. Participants then viewed a bilateral array of colored squares (memory array, 150 ms). The array consisted of 1, 2, 4, or 6 squares with different colors (memory items) in each hemifield (the numbers of squares were always the same between two hemifields). They were presented over two invisible rectangular regions of 3.89 (W) × 6.81 (V), one to the left and another to the right of the fixation point (center-to-center distance: 3.4 deg). A size of each square was 0.73 × 0.73 deg, and its color was randomly chosen from 9 colors (red, green, blue, yellow, magenta, cyan, orange, black, and white). Positions of squares were also randomized on each trial, with the constraint that a distance between squares was 1.94 deg or longer. We asked participants to remember the items in a hemifield indicated by the cue (left or right). After a retention period of 1050 ms, their memory was tested by another array (test array) that was identical to the memory array (no-change trials) or differed by one color (change trials). Participants attended to the cued hemifield and pressed one of two buttons to indicate whether the two arrays were the same (“no-change” response) or not (“change” response). They were also instructed to ignore all items in the uncued hemifield. No time limitation was imposed for the button press.

**Figure 2.**
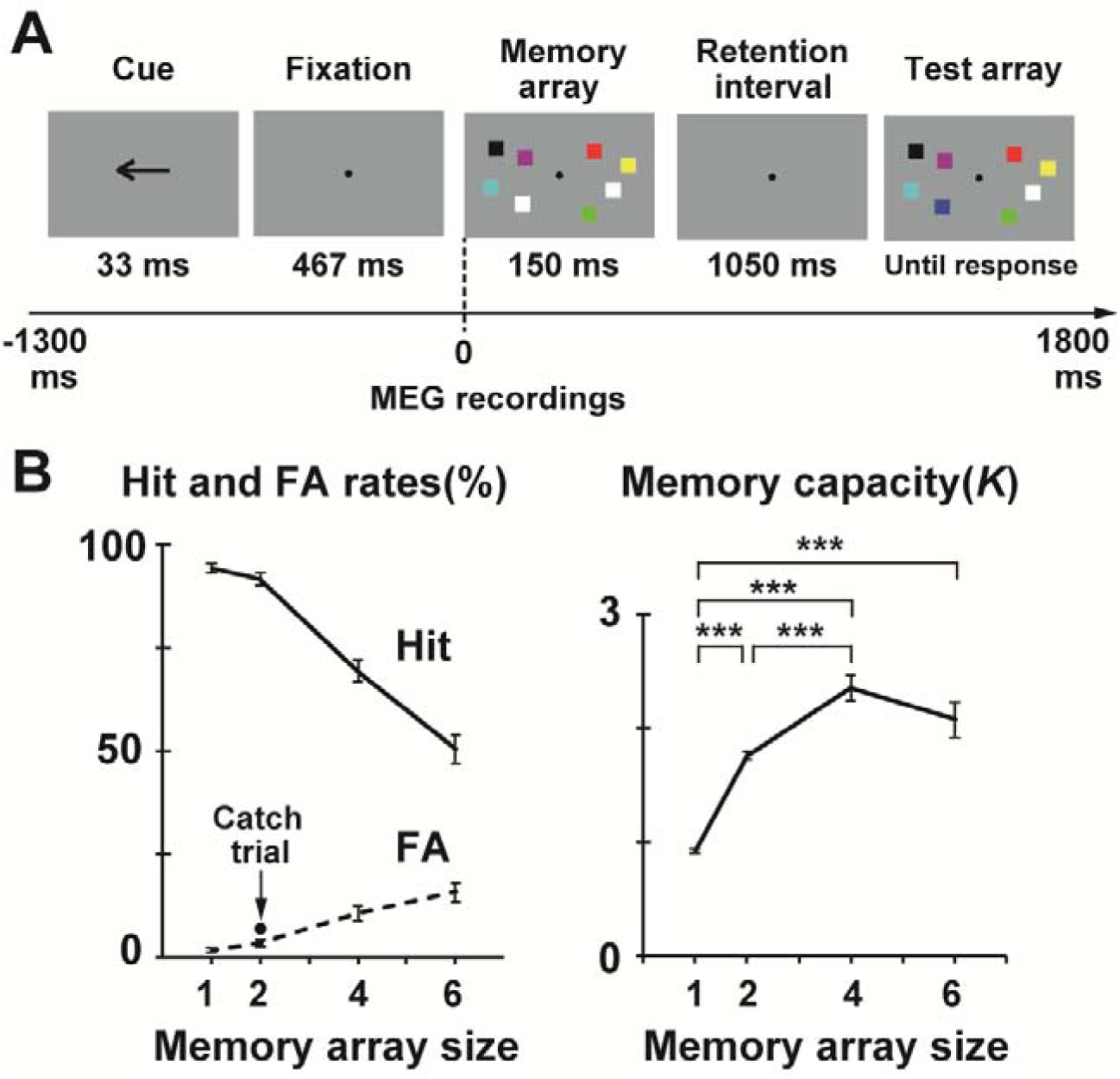
Stimuli and task. (**A**) The cued change-detection task. Participants directed their attention to a cued hemifield (left of right, guided by an arrow at the beginning of each trial) and compared two arrays of colored squares (memory and test arrays) separated by a retention interval. The test array was either identical to the memory array (no-change trials) or differed by one color (change trials). Participants answered whether the two arrays were the same (“no-change” response) or not (“change” response). (**B**) Change in task performance as a function of memory array size (numbers of to-be-memorized items, 1, 2, 4, or 6). A left panel shows hit rates (solid line) in which participants answered “change” in the change trials and false-alarm (FA) rates (dotted line) in which they answered “change” in the no-change trials. A filled black circle indicates a rate of “change” response in catch trials (see texts) where a color change occurred in an uncued hemifield. The right panel shows an index of memory capacity (*K*) estimated from the hit and FA rates (see texts for details). In this and subsequent figures, all error bars denote standard error (SE) across participants. *** *p* < 0.001

A combination of cued hemifield (left or right) and memory array size (WM load, 1, 2, 4, or 6) produced 8 types of trials. We called the trials with leftward cue and 1 memory item as L1, while trials with rightward cue and 6 memory items were named as R6. In addition to those 8 conditions, we introduced 9th condition (catch trials) to check whether participants switched their attention following the cue. In those catch trials, two squares were presented in each hemifield. Although a color change normally took place in the cued hemifield, it occurred in the uncued hemifield in the catch trials (participants thus should press “no-change” button). If participants did *not* allocate their attention following the cue, this would be detected as a high rate of reporting “change” in the catch trials.

An experimental session comprised 100 trials in which 4 catch trials were randomly intermixed with the 96 non-catch trials (L1, L2, L4, L6, R1, R2, R4, and R6, 12 trials for each). A ratio of the change: no-change trials in the non-catch trials was 1:1. Each participant underwent 6 sessions.

Behavioral data were analyzed conforming to previous procedures. For each size of memory array (1, 2, 4, or 6), we computed a hit rate in which participants answered “change” in the change trials and a false-alarm (FA) rate in which they reported “change” in the no-change trials (Fig. 2B, left). Visual memory capacity of each participant (*K*, Fig. 2B, right) was then estimated using a formula below (Pashler, 1988; Cowan, 2001),

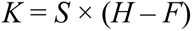

where *K* is the memory capacity, *S* is the size of memory array, *H* and F are the hit and FA rates for that array size, respectively. Previous data showed that the *K* monotonically increased up to memory size 4, reaching a plateau of 2.5 – 3 for higher sizes (Vogel and Machizawa, 2004; Fukuda et al., 2015).

### MEG recordings and preprocessing

Neural activity was measured with a whole-head MEG system (Vector-view, ELEKTA Neuromag, Helsinki, Finland) in National Institute for Physiological Sciences, Okazaki, Japan. This system recorded neural signals at 102 positions over the scalp using 204 planer-type sensors (two sensors at each recording position). One sensor measured changes of neuromagnetic signals in latitudinal directions while the other measured changes in longitudinal directions (sampling rate: 4,000 Hz, analogue band-pass filter: 0.1 – 330 Hz). Data from those two types of sensors were integrated in later analyses (see below). Neuromagnetic waveforms measured by those planar-type gradiometers reflect neural activity in the cerebral cortex just below the recording position.

The preprocessing of MEG data was performed using the Brainstorm toolbox for Matlab (Tadel et al., 2011). Neuromagnetic waveforms were segmented and classified into the 9 conditions above (L1, L2, L4, L6, R1, R2, R4, R6, and catch). An epoch for the segmentation was from −1300 to 1800 ms relative to an onset of a memory array (Fig. 2A). Trials in which a signal variation (a max-min difference within a period of −700 to 1200 ms) was larger than 4,500 fT/cm were excluded from further analyses.

### Inter-peak interval analysis

Speed and regularity of oscillatory signals were evaluated by the IPI analysis. First, raw MEG waveforms were decomposed into 5 frequency bands (delta: 2-4 Hz, theta: 5-7 Hz, alpha/beta: 8-30 Hz, gamma: 31-59 Hz, and high-gamma: 60-100 Hz) with band-pass filters. For each of the 5 filtered waveforms, we measured intervals between contiguous peaks (IPIs). All IPIs during a memory retention period, i.e. 300 - 900 ms from an onset of memory array (Vogel and Machizawa, 2004), were pooled across trials and between latitudinal and longitudinal sensors. This procedure generated a distribution of IPIs (Fig. 1B) at each recording position in each condition. Finally, we computed three parameters of the IPI distribution; mean, standard deviation (SD), and coefficient of variation (CV). The CV was obtained by dividing the SD of IPIs by their mean (CV = SD/mean). It has been used as an index of irregularity (variability) of neural signals (Compte et al., 2003; Qi and Constantinidis, 2015; Standage and Pare, 2018). A higher CV indicates lower regularity of signals.

### Statistical procedures

We first explored the IPI changes related to WM by comparing two types of trials; Memorize-Left trials (L1 – L6) in which participants memorized items in the left hemifield and Memorize-Right trials (R1 – R6) in which they memorized items in the right hemifield. As shown in the left panel of Figure 3B, mean IPIs of 8-30 Hz at right posterior sensors became shorter than those at left posterior sensors in Memorize-Left trials, while retaining items in the right hemifield reduced IPI at the left posterior sensors (middle panel). Contrasting the Memorize-Left and Memorize-Right conditions (right panel) thus isolated hemisphere-specific neural activity related to WM maintenance with total visual inputs in bilateral hemifields equated between conditions. Statistical *t*-maps are shown in Figure 3C. Since a paired *t*-test (N = 24 vs. 24) was repeated for 102 sensor positions, we resolved a problem of multiple comparisons by controlling false discovery rate (FDR). A statistical threshold was adjusted with Benjamini-Hochberg correction (Benjamini and Hochberg, 1995) by setting the *q*-value at 0.05. Sensors showing a significant difference after this correction were marked with white rectangles in Figure 3C.

**Figure 3.**
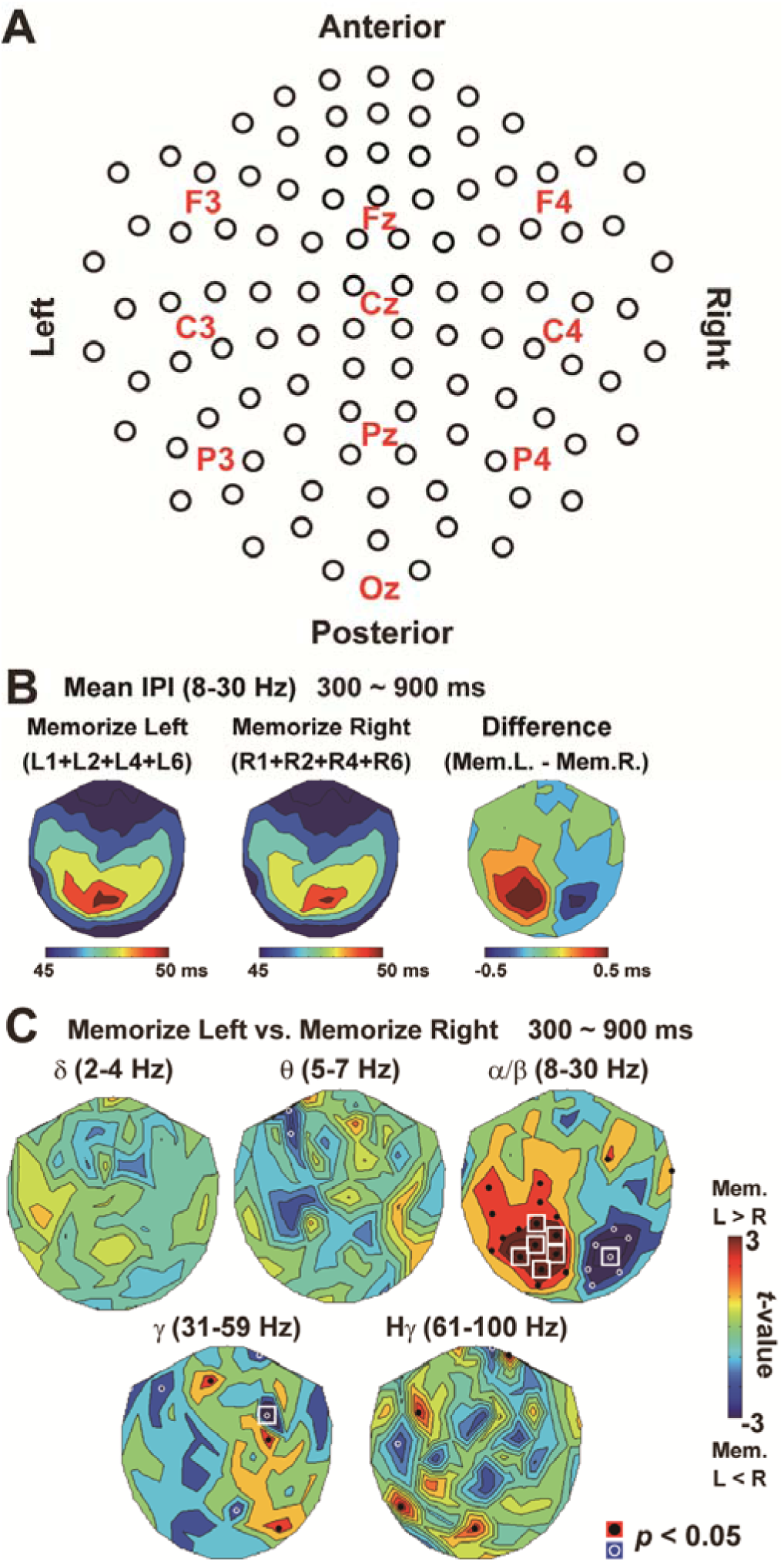
Changes in mean IPIs during the retention period (300 – 900 ms). (**A**) A two-dimensional array of MEG sensors at 102 locations over the scalp. Approximate positions are indicated by labels in the international 10–20 system (e.g. Cz). (**B**) Contour maps of mean IPI over the 102 locations. The left map shows mean IPIs at alpha-to-beta band (8 – 30 Hz) when participants memorized 1-6 items in the left hemifield (L1 – L6), while the middle map shows those when they memorized the right hemifield (R1 – R6). A differential map (L1+L2+L4+L6 vs. R1+R2+R4+R6) in the right panel depicts posterior brain regions showing shorter IPIs when items in contralateral hemifield were retained. (**C**) The *t*-maps of mean IPIs (L1+L2+L4+L6 vs. R1+R2+R4+R6) at five frequency bands from delta (2 – 4 Hz) to high-gamma (61 – 100 Hz). Black dots and white circles denote sensor positions showing a significant (*p* < 0.05) difference. White rectangles indicate a significant difference after a correction of multiple comparisons (see texts). The laterality of retention-related decrease in IPIs was most clearly observed at 8 – 30 Hz.

Another hallmark of memory-related neural activity is a load-dependent changes. We checked this point by contrasting Memorize-Left and Memorize-Right trials for each memory load (L1 vs. R1 and L2 vs. R2, etc. Fig. 4A). If the IPI changes reflected an amount of information stored in memory, the contrast would be clearer as an increase of memory load (Vogel and Machizawa, 2004).

**Figure 4.**
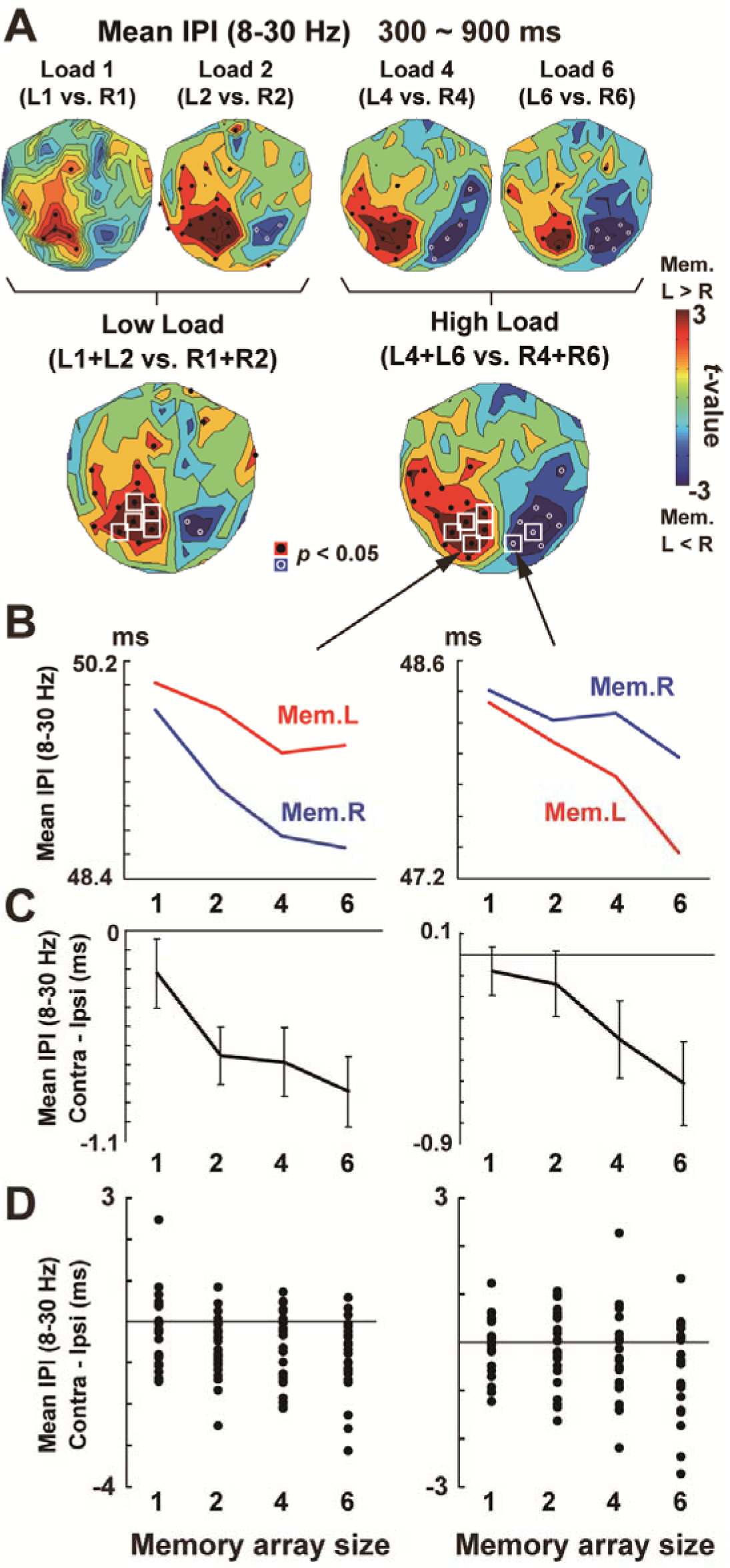
Reduction of mean IPI as a function of memory load. (**A**) A comparison of Memorize-Left vs. Memorize-Right trials in each memory load. The maps in low-load conditions (1 and 2) and high-load conditions (4 and 6) are also shown in the lower panels. (**B**) Mean IPIs at two sensors over the left and right occipito-parietal regions. At both sensors, a two-way ANOVA yielded significant main effects of memory field (left/right) and load (1/2/4/6) as well as their interaction (see texts). (**C**) Differential IPIs. The IPIs became shorter when items in the contralateral hemifield were retained, and this difference grew larger as an increase in memory load. These data indicate that changes in IPIs reflected memory, rather than attention or perception. (**D**) Individual data of the differential IPIs.

## Results

### Behavioral data

Hit and FA rates of the change detection task were shown in Figure 2B. Although the hit rate was above 90 % in array sizes 1 (94.3 ± 1.2 %, mean ± SE across participants) and 2 (91.6 ± 1.4 %), it decreased to 69.4 ± 2.6 % in size 4 and 50.5 ± 3.4 % in size 6. The FA rate, on the other hand, increased monotonically as a function of WM load (size 1: 2.0 ± 0.5 %, size 2: 3.6 ± 0.8 %, size 4: 10.8 ± 1.8 %, size 6: 15.9 ± 2.4 %). Memory capacity *K* (right panel) estimated from those hit and FA rates were 0.92 ± 0.01 (load 1), 1.76 ± 0.04 (load 2), 2.34 ± 0.11 (load 4), and 2.07 ± 0.15 (load 6). A one-way repeated-measures ANOVA with the Greenhouse-Geisser correction on *K* indicated a significant main effect of WM load (*F*(1.45,33.29) = 65.81, *p* < 0.001, *η*^2^ = 0.741). Post-hoc tests with the Bonferroni correction indicated significant difference in loads 1 vs. 2 (*p* < 0.001), 1 vs. 4 (*p* < 0.001), 1 vs. 6 (*p* < 0.001), and 2 vs. 4 (*p* < 0.001).

We also found that a rate of “change” response in the catch trial was much lower (6.9 ± 1.2 %, black dot in **Fig. B**) than the hit rate in load 2 (91.6 %). This result indicates that participants correctly moved their attention into left or right hemifield following the cue direction.

### Comparisons between Memorize-Left and Memorize-Right conditions

Changes in IPIs related to WM maintenance were first explored by a contrast between Memorize-Left (L1-L6) and Memorize-Right (R1-R6) trials. The *t*-maps over the layout of 102 sensor positions (Fig. 3C) revealed that the posterior brain regions showed significant reductions of mean IPIs (8-30 Hz) when visual information in the contralateral hemifield were retained. As shown in Figure 4A, this hemisphere-specific changes in IPIs became more evident in high-load conditions (4 and 6) than low-load conditions (1 and 2). Mean IPIs at two sensors, one over the left and the other over the right occipito-parietal regions, are displayed in Figure 4B-D. At the sensor in the left hemisphere, a two-way ANOVA of memory field (left/right) and WM load (1/2/4/6) indicated significant main effects of memory field (*F*(1,23) = 21.45, *p* < 0.001, *η*^2^ = 0.483) and load (*F*(1.72,39.66) = 8.50, *p* = 0.001, *η*^2^ = 0.270) as well as their interaction (*F*(3,69) = 3.92, *p* = 0.012, *η*^2^ = 0.146). The sensor in the right hemisphere yielded the same results; main effect of memory field: *F*(1,23) = 7.80, *p* = 0.01, *η*^2^ = 0.253, main effect of load: *F*(1.89,43.39) = 10.88, *p* < 0.001, *η*^2^ = 0.321, interaction: *F*(3,69) = 2.95, *p* = 0.039, *η*^2^ = 0.114. Those data showed that the changes in mean IPIs reflected not only attention (main effect of memory field) and perception (main effect of array size) but also an amount of information stored in WM (interaction).

### Comparisons between high-K and low-K individuals

Previous studies reported substantial inter-individual differences in visual WM capacity (Cowan, 2001; Vogel et al., 2001); some people showed high *K* even in high-load memory arrays while others not. Can the IPI changes in alpha/beta band explain those inter-individual differences? We addressed this issue by dividing the 24 participants into two groups based on their *K*s averaged across all conditions (Fig. 5A). When mean IPIs in high-load conditions (L4, L6, R4, and R6) were compared with those in low-load conditions (L1, L2, R1, and R2), the high-*K* participants showed a decreased IPI (high-load < low-load) over a wide area of the brain (Fig. 5B, left). In contrast, this load-dependent reduction was observed only in posterior regions in the low-*K* participants (Fig. 5B, middle). A direct comparison of load effect (high-load minus low-load) between high- and low-*K* individuals (Fig. 5B, right) revealed significant inter-group differences at anterior sensors roughly over the frontal cortex. As shown in Figure 5C, the load-dependent reduction of the frontal IPIs was selectively observed in high-*K* subjects.

**Figure 5.**
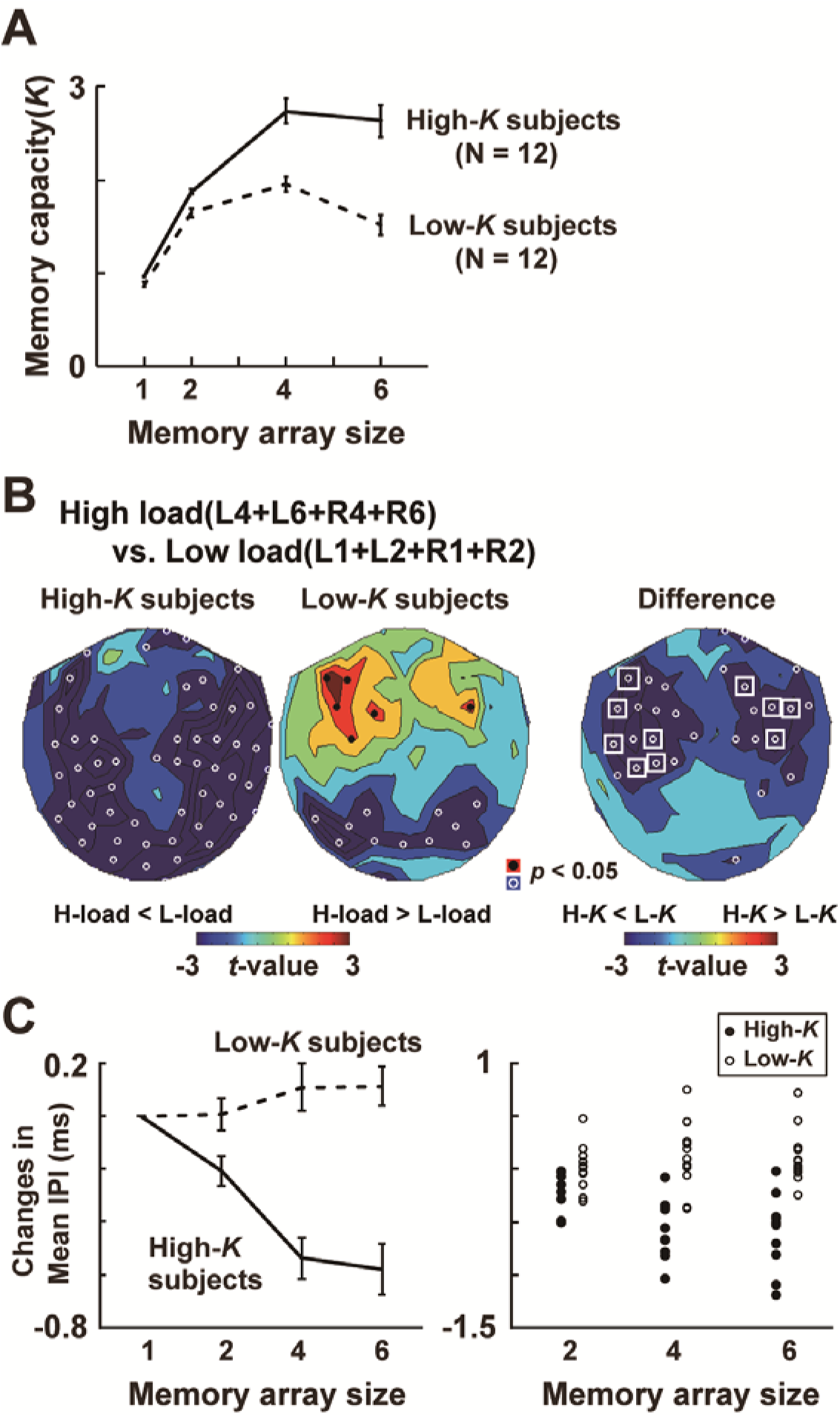
Comparisons of high-*K* and low-*K* participants. (**A**) Memory capacity. The 24 participants were classified into high-*K* or low-*K* groups, based on their mean *K*s across four load levels. (**B**) The *t*-map of high-load (L4+L6+R4+R6) vs. low-load (L1+L2+R1+R2) conditions in high-*K* subjects (left panel) and low-*K* subjects (middle panel). Sensor positions showing shorter IPIs in high-than low-load conditions are shown in blue. In the right panel, differential IPIs (high-load minus low-load) are compared between high-*K* and low-*K* subjects (inter-group difference). High-*K* subjects were characterized by a load-dependent reduction of mean IPIs in anterior brain regions. (**C**) Changes in IPIs (left: mean ± SE, right: individual data) averaged across 10 sensors marked by white rectangles (*p* < 0.05, corrected) in panel **B**. Relative changes from load 1 condition are shown.

Interestingly, the difference between high-*K* and low-*K* groups was also seen in IPIs during a baseline period, but in an opposite direction. Figure 6A displays the contour maps of IPIs in the baseline period (pre-cue period, from −1300 to −700 ms). The mean IPIs were longer in high-*K* than low-*K* individuals, indicating that higher task performance was associated with slower brain rhythms over the frontal cortex. When faced with high-load displays, high-*K* subjects memorized a large number of items by making their slow rhythm faster (Fig. 6B). These data suggest that a key feature of high-*K* individuals was the changeability (flexibility) of brain rhythms, not a speed (absolute velocity) of oscillatory signals.

**Figure 6.**
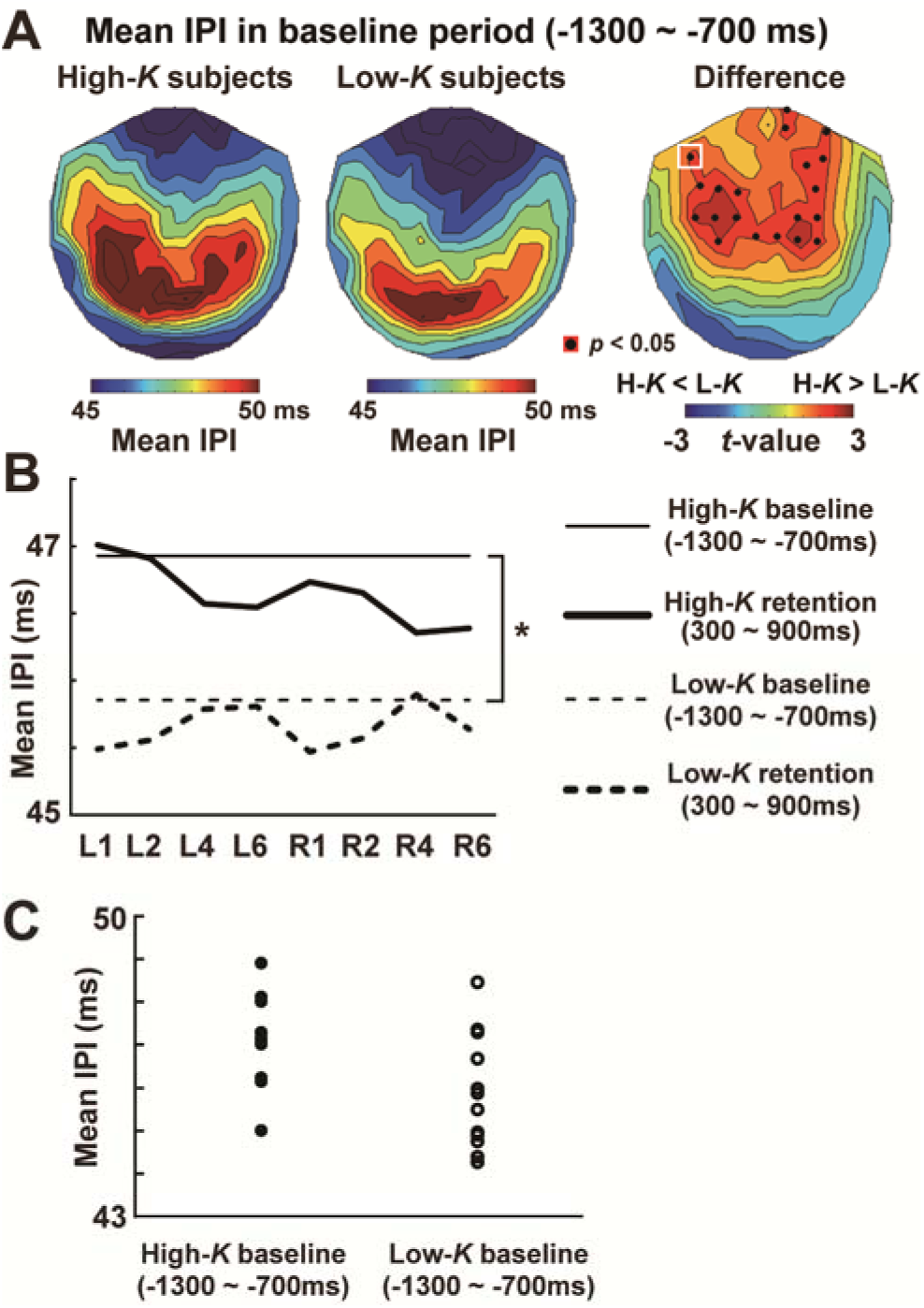
Mean IPIs at 8 – 30 Hz during a baseline period (from −1300 to −700 ms). (**A**) Contour maps of baseline IPIs in high-*K* subjects (left), low-*K* subjects (middle), and an inter-group difference (*t*-map). The baseline IPIs were obtained as a mean across 8 conditions from L1 to R6. (**B**) IPIs in the retention (thick lines) and baseline (thin lines) periods of high-*K* (solid lines) and low-*K* subjects (dotted lines) at the left anterior sensor (white rectangle in panel **A**). High-*K* subjects were characterized by slower alpha-to-beta rhythm in the baseline period. When faced with high-load displays, they memorized a large number of items by making the slow rhythm faster. These data suggest that larger capacity was associated with greater changeability (flexibility) of brain rhythms. * *p* < 0.05 (**C**) Individual data of the baseline IPIs in high-*K* (filled dots) and low-*K* (open dots) subjects.

### Memory-related changes in a variance of IPIs

We finally analyzed changes in variance of IPIs related to vWM. Figure 7 shows the CV (variance) of alpha/beta IPIs at 300 – 900 ms compared between Memorize-Left and Memorize-Right conditions. Smaller CVs denotes higher regularity of oscillatory signals. The CVs at right-hemisphere sensors became smaller when participant retained items in the left hemifield (colored in blue), whereas memorizing right-hemifield items reduced the variance at sensors over the left hemisphere (red). This pattern was most clearly observed in the low-load conditions of high-*K* participants.

**Figure 7.**
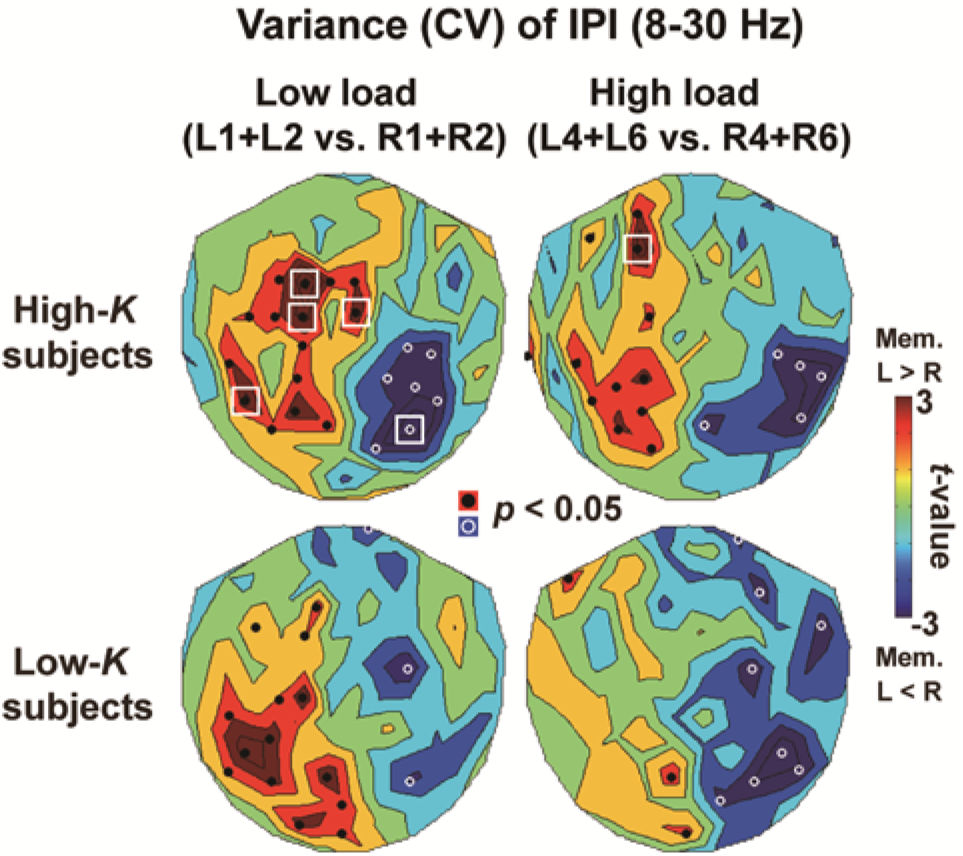
Variance of IPIs during the retention period (300 – 900 ms). We computed the coefficients of variation (CVs), which was a standard deviation of IPIs divided by their mean (CV = SD/mean). Smaller CVs denotes smaller variance of IPIs (high regularity of oscillatory signals). Comparisons between Memorize-Left and Memorize-Right conditions reveals reduced CVs at sensors contralateral to a memory field, especially when high-*K* participants retained the information of low-load arrays (upper left).

## Discussion

In the present study, we investigated changes in brain rhythms related to a retention of vWM. We found a load-dependent decrease of mean IPIs in the occipito-parietal regions contralateral to a memory field. A comparison between high-*K* and low-*K* participants further indicated a critical role of the anterior (frontal) regions in memorizing a large number of items simultaneously.

### Reduction of mean IPIs and WM maintenance

Several studies have investigated a relationship between vWM and a speed of oscillatory signals in the fontal and parietal cortex. No consistent results, however, have been obtained so far. Using the Sternberg WM task, Babu Henry Samuel et al. (2018) measured a peak frequency of alpha rhythm (8-12 Hz) during a retention period of a digit set (e.g. 59713). They showed a negative correlation between oscillation frequency and behavioral data; trials with higher alpha frequency (faster rhythm) was associated with lower performance of WM task (Babu Henry Samuel et al., 2018). Consistent with this view, Cohen (2011) investigated a peak frequency at 2-18 Hz in a retention period of a visual stimulus (line drawing), reporting that individuals with higher peak frequency showed lower performance in memory task (Cohen, 2011). In contrast, Moran et al. (2010) used the change detection task and reported that the peak frequency at 4-12 Hz was significantly higher in high capacity individuals than low capacity individuals (Moran et al., 2010). According to Moran et al. (2010), their data were consistent with the communication theory (Shannon, 1948) predicting a higher capacity of information transfer achieved by an increase in peak frequency.

Although it is unclear what produced the mixed results in a previous literature, a hallmark of the present study was a measurement of IPIs at a wider bandwidth over alpha-to-beta range (8-30 Hz). This approach is based on a previous view that alpha and beta rhythms play a similar role in WM and thus can be seen as a unified band of oscillatory signals (Lundqvist et al., 2011; Roux and Uhlhaas, 2014; Miller et al., 2018). The mean IPIs identified with a pass band of 8-30 Hz actually showed clear decrease along with an increase in memory load (Fig. 4), which was associated with a better performance of WM task (Fig. 5). These results suggest that neural temporal codes related to WM are distributed in a broad frequency range over alpha to beta bands.

### Inter-individual differences of mean IPIs in a baseline period

A comparison of high-*K* and low-*K* groups (Fig. 5) revealed that the high-*K* individuals showed a significant reduction of frontal IPIs in high-load conditions (L4, L6, R4, and R6) whereas the low-*K* individuals did not. It should be noted, however, that those results do *not* indicate a direct link between a faster brain rhythm and larger memory capacity, because high-*K* subjects showed a slower brain rhythm in a baseline period (Fig. 6). A key feature of high-*K* subjects therefore was the greater flexibility (changeability) of brain rhythms from low-load to high-load conditions. In previous studies, a successful maintenance of WM was associated with a high rate of neuronal firing in the prefrontal cortex (Constantinidis et al., 2018) or a formation of new patterns of brain rhythm, such as the cross-frequency coupling between theta and gamma waves (Lisman and Jensen, 2013). Our study proposes a new possibility that multi-item WM is achieved by a flexible modulation of ongoing brain rhythms in accordance with memory loads.

### Reduced variance of IPIs in a retention period

We also found a significant decrease in variance of IPIs (increase in regularity of brain rhythms) during a retention interval (Fig. 7). Although several studies on human WM have analyzed regularity of neural activity using correlation dimension (Grassberger and Procaccia, 1983), approximate entropy (Pincus, 1991) and Hurst exponent (Hurst, 1951), no consistent result has been obtained (Zarjam et al., 2012; Behzadfar et al., 2017). We presently approached this issue using the IPI analysis and found the memory-related increase in regularity. Our results support the attractor hypothesis in WM, because such attractor states are thought to emerge from self-sustained recurrent networks that can produce a periodic (regular) oscillatory signal.

Different from mean IPIs, the changes in regularity was more clearly observed in low-than high-load conditions (Fig. 7). Our behavioral data (Fig. 2B) showed that those low-load trials were characterized by higher hit rates and lower FA rates than high-load trials. The increased regularity of brain rhythm therefore might reflect a high fidelity of WM contents in those low-load trials.

## Acknowledgments

This work was supported by KAKENHI Grants Number 19H04430 from the Japan Society for the Promotion of Science (JSPS) to Y.N. We thank Mr. Y. Takeshima (National Institute for Physiological Sciences, Japan) for his technical supports. The authors declare no competing financial interest. All data supporting the findings of this study are available from Y.N. upon reasonable request.

